# Microglia Mitochondria Drive Neuronal Maturation via Metabolic and Transcriptional Reprogramming

**DOI:** 10.1101/2025.04.29.651306

**Authors:** Sydney P. Sterben, Charita C Anamala, Vaishnavi Koduri, Sahan B.S. Kansakar, Volha Liaudanskaya

## Abstract

Autism Spectrum Disorder (ASD) is a complex neurodevelopmental condition characterized by impaired social interactions, repetitive behaviors, and disrupted neuronal circuit maturation. Emerging evidence implicates both microglial function and mitochondrial regulation as critical determinants of ASD pathogenesis. Here, we identified microglia and their mitochondria as active modulators of neuronal circuit development, highlighting their potential roles as mechanistic contributors and biomarkers in ASD progression.

**Graphical Abstract:** 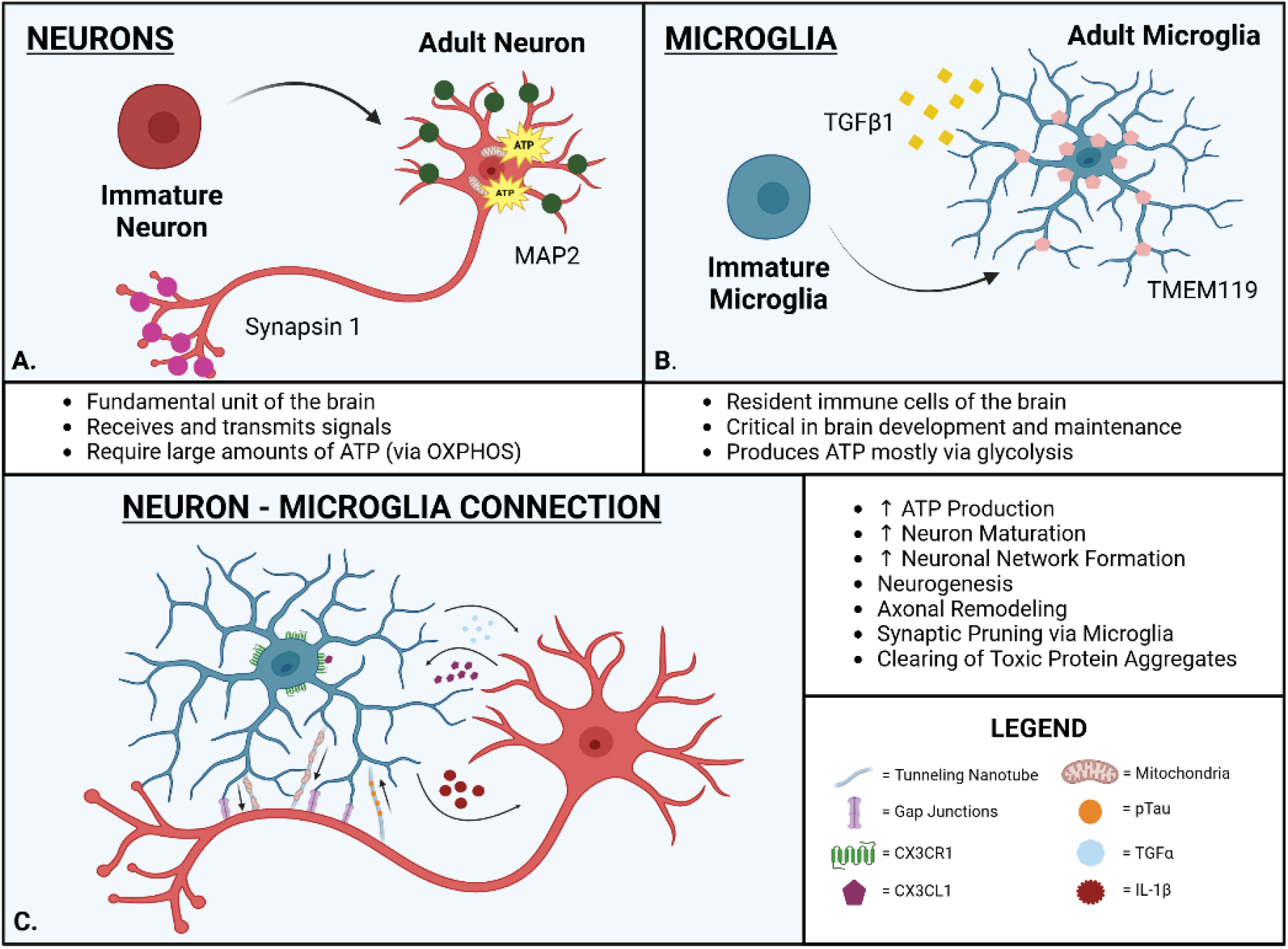

## Introduction

The development and maturation of neurons is a well-orchestrated and highly dynamic process that guides functional neural circuit formation (1). This process extends from early neurogenesis through adolescence and adulthood, particularly in the human brain, which undergoes significant alterations after birth. Neuronal maturation is regulated by a complex interplay of intrinsic genetic programs and extrinsic environmental signals, including neuron-glia interactions, synaptic activity, secreted molecules, metabolic state, and mitochondrial function (2–4). Any disruption in these developmental programs can lead to broad alterations in neuronal architecture and function, manifesting in abnormal synapse formation, delayed neurite outgrowth, impaired GABAergic signaling, increased oxidative stress, and neuroinflammation (5).

Autism spectrum disorder (ASD) is a neurodevelopmental condition increasingly associated with disrupted neuronal maturation (6)(7). Affecting approximately 1 in 36 children in the United States (8), ASD is characterized by deficits in social interaction and communication, along with restrictive and repetitive behaviors (9). While advances in diagnostic criteria and awareness have contributed to the rise in ASD diagnoses, the disorder remains diagnosed solely through behavioral observation (10, 11). This reflects the persistent lack of molecular biomarkers and an incomplete understanding of its underlying biology (9). Moreover, comorbidities such as epilepsy, gastrointestinal disturbances (6, 12), and sensory processing disorders often exacerbate the clinical burden and diminish the quality of life. Existing therapies are largely symptomatic, targeting irritability and behavioral issues rather than the root causes of the disorder, underscoring the urgent need for biologically informed treatment strategies (7).

Neurodevelopmental models increasingly implicate impaired neuronal maturation and aberrant synaptic pruning as contributing factors in ASD etiology (13–15). During early postnatal development, the brain generates an abundance of synaptic connections, which are subsequently refined through activity-dependent pruning to establish efficient and adaptable neural circuits (16). Microglia—the brain’s resident immune cells—are central to this process (13). They regulate the elimination of excess synapses, modulate dendritic remodeling, and help balance excitatory and inhibitory signaling within the CNS (17). Dysregulation of microglial activity, including impaired synaptic engulfment and disrupted neuron-microglia signaling (e.g., via the CX3CL1–CX3CR1 axis), has been implicated in several ASD models and is associated with behavioral phenotypes characteristic of the disorder (18). These findings emphasize the critical importance of microglial-neuronal communication during early brain development.

At the molecular level, disruptions in the expression of neuronal maturation markers— such as Tuj1, MAP2, and synapsin-1—along with abnormal microglial phenotypes marked by altered TMEM119 and TGFβ1 expression, have been observed in ASD patient tissue and animal models (19, 20). Together, these data point to a broader failure of neuro-glial crosstalk, contributing to the developmental deviations observed in ASD patients.

In parallel with defective pruning, mitochondrial dysfunction has emerged as a key contributor to the pathogenesis of ASD. Beyond their role in cellular energy production, mitochondria are now recognized as central regulators of neuronal proliferation, differentiation, and synapse formation (21), (22). The high metabolic demands of the developing brain require efficient mitochondrial oxidative phosphorylation (OXPHOS) (23, 24). Yet, individuals with ASD often exhibit impaired mitochondrial function, as evidenced by reduced ATP levels, dysregulated electron transport chain (ETC) activity, elevated oxidative stress, and abnormal metabolite profiles—such as increased lactate and pyruvate levels (25–27). Further, disruptions in mitochondrial dynamics, including an imbalance between fission and fusion (28) (29), lead to fragmented mitochondria, ROS accumulation, and neuronal dysfunction—hallmarks also observed in postmortem ASD brain tissue (30, 31).

Microglia-neuron metabolic crosstalk has recently emerged as a novel mechanistic axis implicated in neurodevelopmental disorders. Microglia are capable of transferring functional mitochondria to stressed neurons, modulating metabolic profiles, and enhancing mitochondrial plasticity. These findings suggest that microglia contribute to neuronal development through synaptic remodeling and directly influence neurons’ bioenergetic capacity. The precise mechanism of this support, whether through physical contact, secreted vesicles, or metabolic intermediates, remains poorly understood, particularly in the context of neurodevelopment.

Despite growing mechanistic insights, it remains unclear whether physical cell-cell contact between microglia and neurons is required for these beneficial effects or whether microglial-derived mitochondria and metabolites can independently influence neuronal maturation. To address this knowledge gap, we utilized a human 3D in vitro model of the brain (32) to investigate the role of microglial mitochondrial support in neuronal development. Specifically, we evaluated (1) whether direct microglia-neuron interaction is necessary for promoting neuronal maturation, (2) the functional impact of mitochondrial transfer on neuronal bioenergetics, and (3) how our model recapitulates in vivo-like neurodevelopmental features. We found that treatment with microglial mitochondria partially promotes neuronal maturation and induces transcriptional alterations as early as 48 hours post-treatment. However, factors secreted by microglia in combination with a physical connection between neurons and microglia are necessary for complete development. In summary, our study provides insight into the mechanistic basis of microglial support during neuronal maturation. It offers a platform for identifying new therapeutic strategies targeting cellular dysfunction in neurodevelopmental disorders.

## Results

### Microglia promote neuronal maturation and network development in a 3D human in vitro model

To explore the benefits of microglia in promoting neuronal maturation, we used our 3D *in vitro* human tissue model composed of induced neuronal stem cells (iNSCs) and an HMC3 microglia cell line with fluorescently tagged mitochondria (dsRED2 and EBFP2, respectively) enabling direct comparison of Neuron only monocultures (N) and Neuron + Microglia (NM) co-cultures to naïve neuron monocultures (Figure 1A) (41, 42). We analyzed the presence of neuron-specific maturation markers at 2wks and 5wks post-seeding to demonstrate the developmental differences between the two experimental conditions.

**Figure 1.**
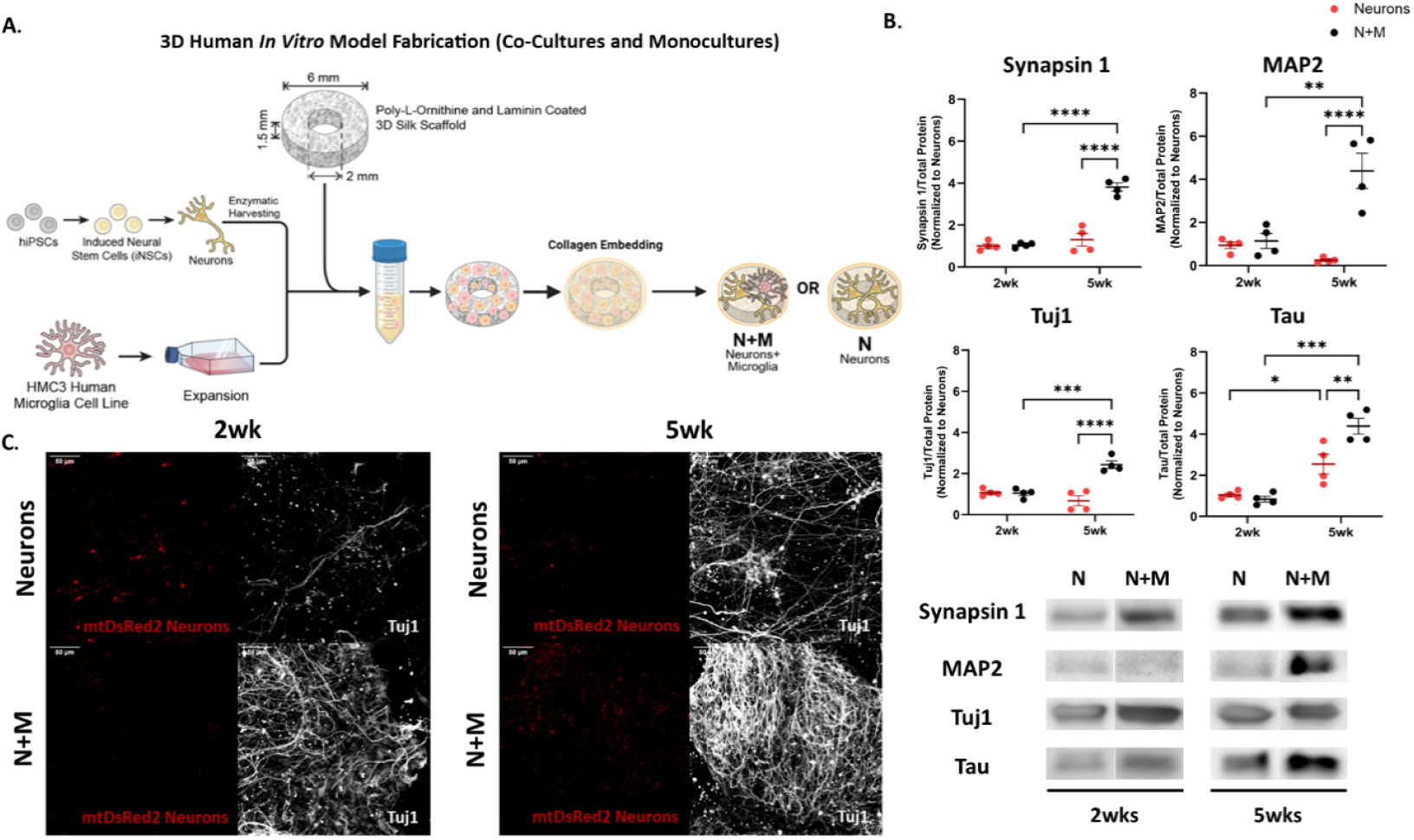
Microglia boosts Neuronal Maturation – **A**. Schematic representation of the preparation of our NM co-culture and neuron monoculture. **B**. Western blot quantification of select neuron maturation markers (Synapsin 1, MAP2, Tuj1, and TAU) with Neurons in red and NM in Black. n = 4 for NM and Neurons. Two-way ANOVA looking at effects of time and culture type with Tukey post-hoc test and α = 0.05. ROUT outlier analysis method. Analysis completed in GraphPad Prism. **C**. Representative images of Tuj1 and mtDsRed2 neuron mitochondria. Nikon A1 inverted LUNV confocal microscope 20x objective 1024×1024 pixels with 50-steps Max projection. Scale bar: 50µm.

At 2 weeks post-seeding, we observed no statistical difference in the expression of Synapsin 1, MAP2, Tau, or Tuj1 between neurons and NM conditions (p-value > 0.05). In contrast, by 5wks, NM co-cultures exhibited significant upregulation of the above-mentioned maturation markers compared to neurons (p-value < 0.0001 for Synapsin 1, MAP2, and Tuj1, p-value < 0.01 for Tau). These results suggest accelerated and more robust neuronal maturation in the presence of microglia.

Immunofluorescence analysis further confirmed enhanced Tuj1 expression and extensive neuronal network formation in NM cultures at both time points, with pronounced network complexity evident by 5 weeks (Figure 1C). While network structures were present in neuron-only cultures, NM co-cultures showed greater neurite density and connectivity, suggesting more advanced functional integration. At last, the interconnectivity of the mitochondrial network also visually increased at 5wks in the NM cultures compared to the naïve neuron condition. Mitochondrial networks are crucial in the functionality of cells like neurons that rely on high levels of ATP. Together, these results support a critical role for microglia in orchestrating neuronal maturation and network assembly, potentially through both trophic and metabolic mechanisms.

### Microglia Mitochondria Reprogram Neuronal Bioenergetics in a 3D Human Brain Model

To determine the contribution of microglial mitochondria in neuronal bioenergetic maturation, we isolated fresh microglial mitochondria from 3D microglia monocultures and applied them directly to naïve neuronal cultures. Bioenergetic profiling was performed at 24hrs, 48hrs, and 1wk post-treatment using Seahorse XF analysis to measure Oxygen Consumption Rate [OCR] and Extracellular Acidification Rate [ECAR]) (Figure 2A and B).

**Figure 2.**
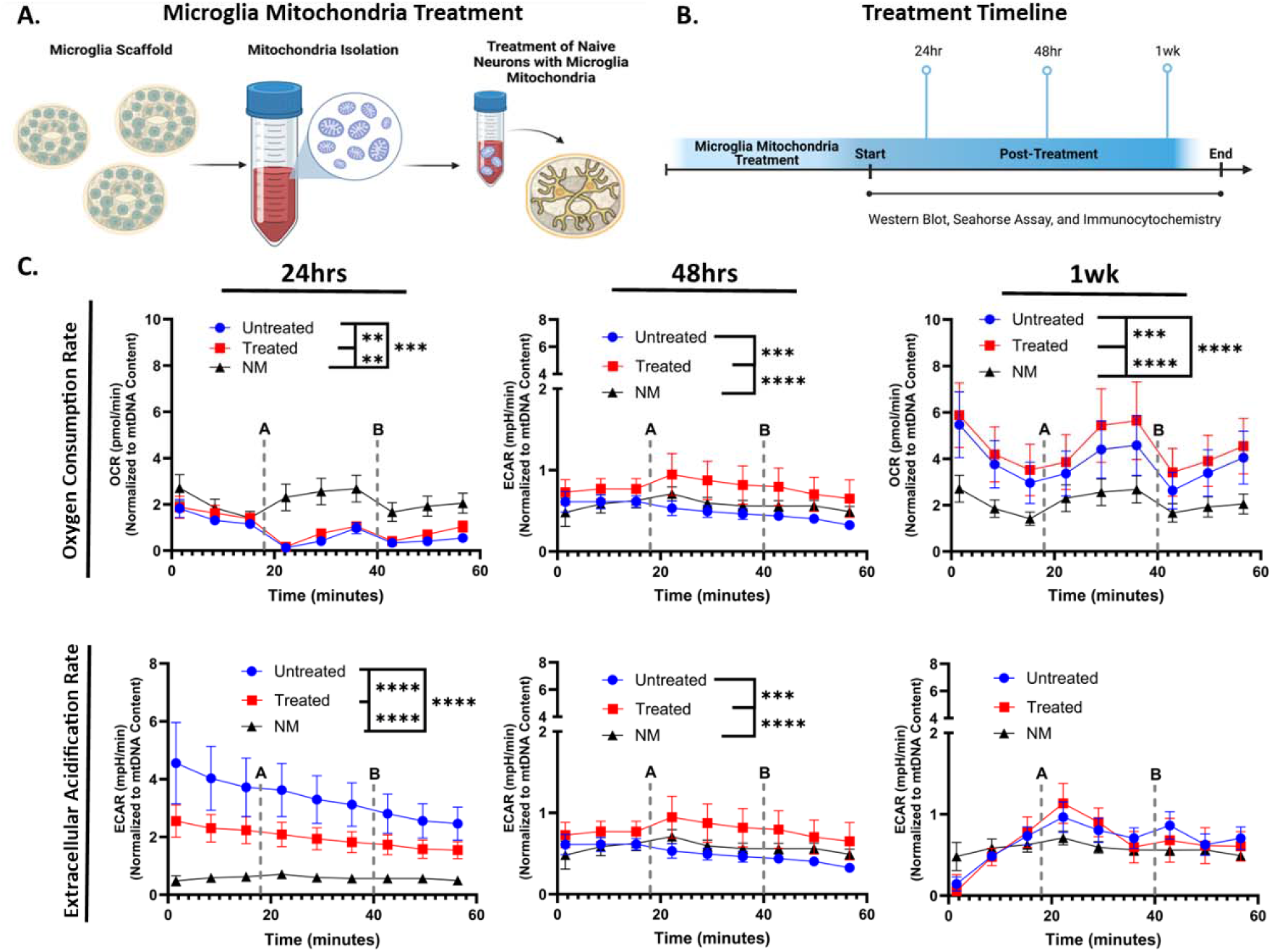
Mitochondria Enhance Neuronal Bioenergetics over time. **A**. Schematic representation of the experimental design. **B**. Treatment and analysis timeline. **C**. Real-Time ATP Rate Assay results comparing OCR and ECAR between different conditions across select time points. n = 3 scaffolds NM and Untreated. n = 4 for Treated. Mean ± SEM. One-way repeated measures ANOVA with Tukey post-hoc test and α = 0.05. ROUT outlier analysis method. Analysis completed in GraphPad Prism. Each timepoint was replicated at least twice.

Mitochondria transfer induced significant alterations in neuronal metabolic activity. At 24 hours, OCR was significantly elevated in treated neurons compared to NM (p<0.01), and untreated neurons (p<0.001), indicating a rapid enhancement in oxidative phosphorylation. This trend persisted at 48hrs, where OCR in the treated group remained comparable to NM (p-value > 0.05), suggesting metabolic stabilization. By 1wk, the Treated condition exhibited the highest OXPHOS activity across conditions (p-value < 0.0001 vs NM, p < 0.001 vs untreated), while untreated neurons also showed increased OCR relative to NM (p < 0.0001), likely reflecting compensatory stress responses.

ECAR analysis further supported a metabolic shift. At 24 hours, untreated neurons displayed elevated ECAR compared to both treated and NM (p<0.0001), indicative of heightened glycolysis. However, by 48 hours, ECAR significantly increased in treated neurons (p < 0.001 vs. Untreated; p < 0.0001 vs. NM), aligning with enhanced glycolytic activity. These effects resolved by 1 week, with no significant differences in ECAR among groups.

Collectively, these findings demonstrate that treatment with microglial mitochondria can modulate neuronal bioenergetics, enhancing both oxidative and glycolytic capacity in a temporally dynamic manner. This suggests that microglial mitochondrial transfer may be a key regulatory mechanism in neuronal metabolic maturation.

### Microglial Mitochondria Restore Neuronal Mitochondrial Dynamics

In the NM co-cultures, microglial and neuronal mitochondria show clear colocalization (Figure 3A). While there were alterations on a bioenergetic level at 24hrs, there were no significant alterations to protein expression or mitochondrial health at said timepoint. Due to this, the 24hr timepoint was removed from analysis (Figure S1). By 48hrs post-treatment, we observed visual confirmation of microglia mitochondria integrating into neurons, a trend that persisted through the 1wk time point, supporting the ability of neurons to internalize extracellular mitochondria for potential metabolic support (32). Spearman’s correlation coefficients for colocalization of microglial mitochondria did not reach statistical significance; a modest upward trend was observed from 48 hours through 1 week (Figure 3B).

**Figure 3.**
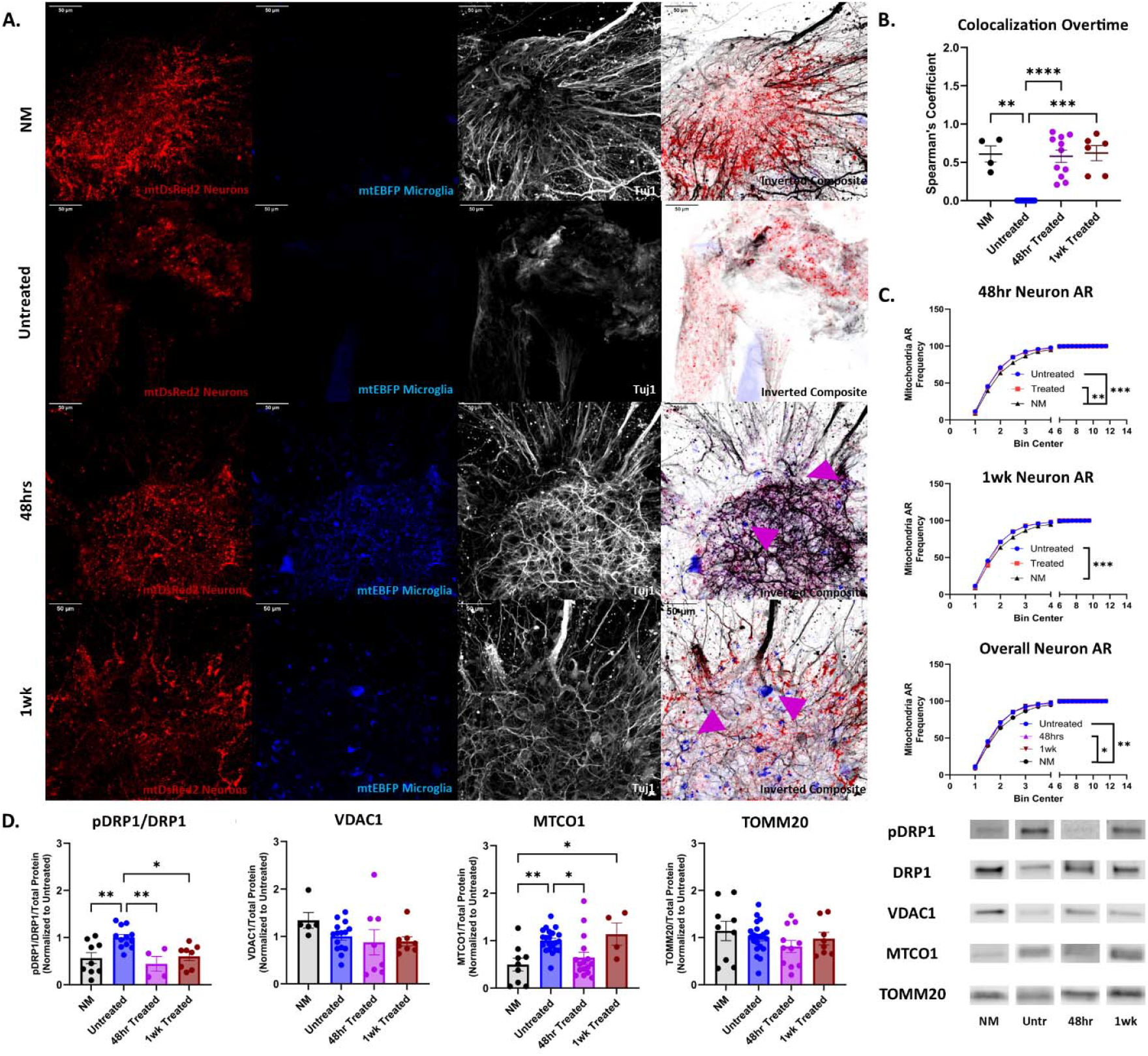
Mitochondrial Health and Colocalization: **A**. Representative images of NM, Untreated, 48hrs Treated, and 1wk Treated. mtDsRed2 neurons and mtEBFP2 microglia. Tuj1 staining for neuronal networks. Nikon A1 inverted LUNV confocal microscope, 20x objective, 1024×1024 pixels, with 50-steps Max projection. Scale bar: 50µm. **B**. Colocalization analysis was performed with JaCoP using Spearman’s Correlation Coefficient comparing the colocalization of mtDsRed2-positive neuronal mitochondria and mtEBFP-positive microglial mitochondria. **C**. Mitochondria aspect ratio comparing the length and width of mitochondria at select time points (48hr and 1wk) along with NM and Untreated. **D**. Mitochondria health markers, pDRP1/DRP1, TOMM20, VDAC1, and MTCO1 intracellular mitochondria western blots normalized to total protein concentration per lane and to the Untreated condition. n = 3 for NM and Untreated. n = 4 for 48hr and 1wk Treated. Mean ± SEM. One-way ANOVA with Tukey post-hoc test and α = 0.05. ROUT outlier analysis method. Analysis completed in GraphPad Prism. Each timepoint was replicated at least twice.

Quantitative analysis of mitochondrial morphology revealed that, at 48 hours, neurons treated with microglial mitochondria exhibited significantly more fragmented mitochondria (reduced aspect ratio [AR]) compared to NM (p < 0.01), consistent with a fragmented mitochondrial state. However, by 1-week post-treatment, AR values in treated neurons no longer differ from NM, indicating a restoration of mitochondrial morphology. In contrast, untreated neurons retained a significantly lower AR compared to both 48-hour and 1-week treated groups (p < 0.001), suggesting persistent mitochondrial fragmentation in the absence of treatment (Figure 3C).

Lastly, to assess mitochondrial health, we examined markers of fission and function. Phosphorylation of DRP1 at serine 616 (a key marker of mitochondrial fission) was significantly elevated in untreated neurons compared to NM (p < 0.01), 48-hour treated (p < 0.01), and 1-week treated groups (p < 0.05), supporting the AR findings (Figure 3D). While TOMM20 and VDAC1 levels remained unchanged across conditions, intracellular MTCO1 expression was highest in untreated neurons (vs. NM, p < 0.01; vs. 48-hour treated, p < 0.05). Interestingly, 1-week treated neurons showed increased MTCO1 compared to NM (p < 0.05), suggesting potential compensatory mitochondrial activity at later timepoints. Together, these data suggest that treatment with microglial mitochondria promotes uptake into neurons and supports improved mitochondrial dynamics, specifically by reducing fission and restoring healthier mitochondrial morphology.

### Microglial Presence Enhances Neurodevelopment Beyond Mitochondrial Transfer

To investigate whether early bioenergetic changes induced by microglial mitochondria are accompanied by transcriptional reprogramming, we performed RNA sequencing on neurons treated for 48 hours with isolated microglial mitochondria compared to untreated controls. Differential gene expression analysis revealed significant enrichment of nervous system development pathways, including neurogenesis (FDR = 0.0041), nervous system development (FDR = 0.0033), and neuron differentiation (FDR = 0.0198), all with adjusted p-values < 0.05 and log_2_(fold change) ≥ 0.2 (Figure 4A). Upregulated genes included ID1, ID3, and SALL4 (regulators of proliferation and differentiation), as well as RUNX1, LAMB1, PLXND1, and MED12, which are associated with neuronal development. These transcriptional changes at 48 hours preceded observable phenotypic differences at 1-week post-treatment, as evidenced by increased MAP2 expression—a marker of dendritic maturation—in treated neurons compared to untreated controls (p < 0.0001). To confirm the absence of neurodegenerative effects, we compared Tuj1-positive neural networks and the presence of key neurodegenerative markers (amyloid precursor protein [APP], transactive response DNA-binding protein 43 [TDP43], and phosphorylated tubulin associated unit [pTau]) to the NM co-culture. The treatment did not adversely affect neurodegenerative markers or neuronal networks (Figure S2).

**Figure 4.**
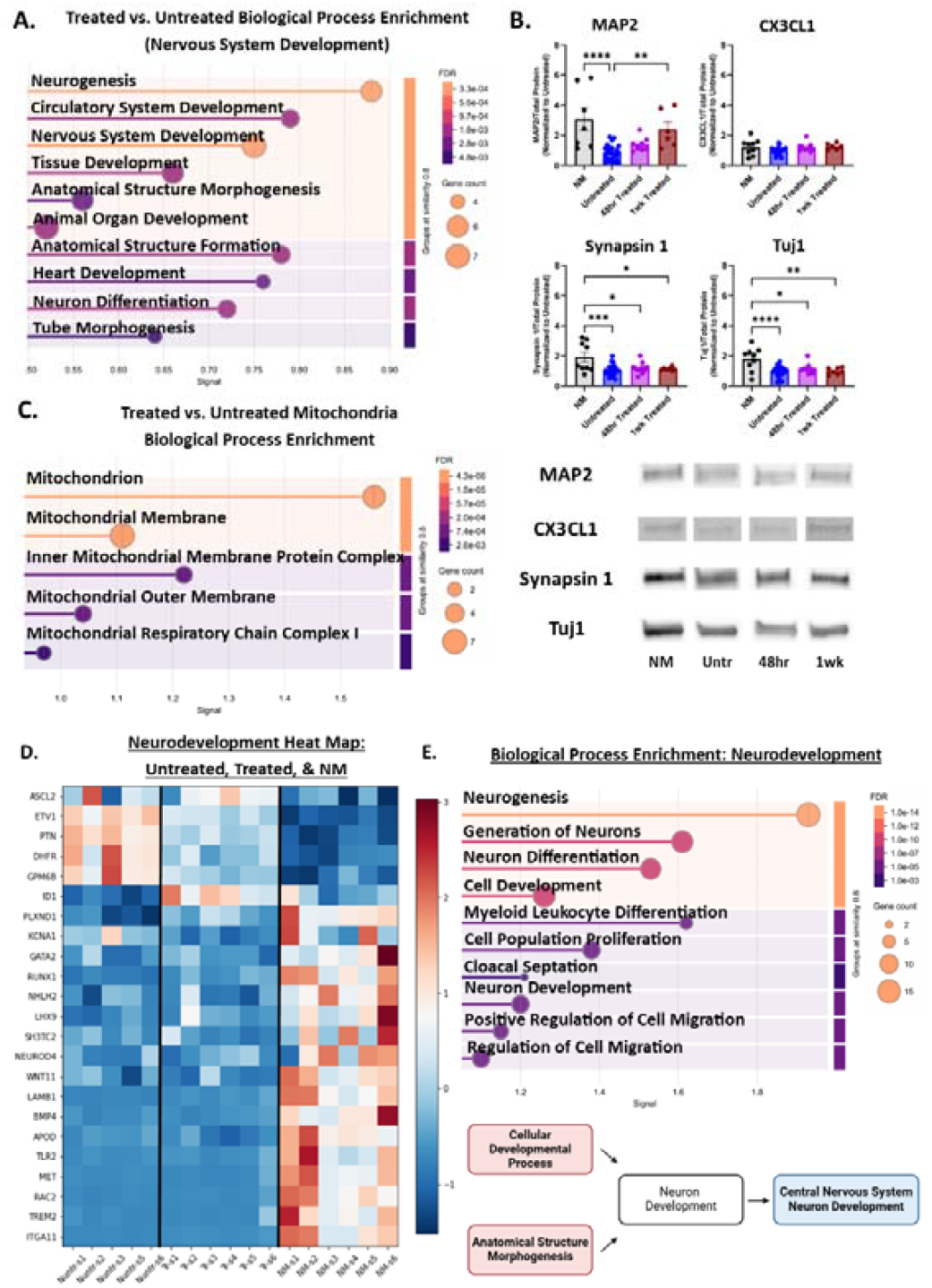
Treatment Effect on Nervous System Development and Mitochondria: **A**. Biological process enrichment, specifically targeting nervous system development between treated and untreated. Due to wanting to see the finer details in the differences between the treated and untreated conditions, the Log_2_(Fold Change) threshold was set to 0.2 for analysis. However, an adjusted p-value of < 0.05 was still required to be deemed “significant”. Medium confidence interaction score (0.4). False Discovery Rate (FDR) ≤ 0.05. **B**. Neuronal maturation markers MAP2, CX3CL1, Synapsin 1, and Tuj1 over time western blots normalized to total protein concentration per lane and to Untreated condition. **C**. Biological process enrichment, specifically mitochondria-related genes from Mito Carta 3.0 between treated and untreated. Log_2_(Fold Change) = 0.2, adjusted p-value of < 0.05, medium confidence interaction score (0.4), and FDR ≤ 0.05 (33). **D**. Comparison Between NM vs. Treated Heat map of z-score normalized DEGs comparing NM with 48hr treated scaffolds. P-value < 0.05 required to be deemed significant for further analysis. Medium confidence interaction score (0.4). False Discovery Rate (FDR) ≤ 0.05. **E**. Biological process enrichment focusing on neurogenesis between NM and 48hr Treated. Log_2_(Fold Change) = 1.0, adjusted p-value of < 0.05, medium confidence interaction score (0.4), and FDR ≤ 0.05. Mean ± SEM. One-way ANOVA with Tukey post-hoc test and α = 0.05. ROUT outlier analysis method. Analysis completed in GraphPad Prism. n = 3 for NM and Untreated. N = 4 for 48hr and 1wk Treated. Each timepoint was replicated at least twice.

Focusing on mitochondrial-specific genes from the MitoCarta 3.0 database, we identified seven genes significantly altered by treatment. These genes were enriched in pathways related to the “inner mitochondrial membrane protein complex” (FDR = 0.0062) and “mitochondrial respiratory chain complex I” (FDR = 0.0265) and were predominantly downregulated in treated neurons. The only upregulated mitochondrial gene was CPT1B, a key regulator of fatty acid oxidation. Despite downregulation of ETC-related genes, neurons treated with microglial mitochondria demonstrated increased oxidative metabolism (Figure 2C), suggesting a compensatory shift toward fatty acid oxidation. These results suggest that while neurons incorporate extracellular mitochondria, this may be accompanied by transcriptional remodeling that attenuates endogenous ETC gene expression.

To determine whether mitochondrial transfer alone could replicate the effects of direct microglia-neuron interaction, we compared the transcriptional profiles of the 48-hour Treated condition to the NM co-culture. Although mitochondrial treatment promoted neurodevelopmental gene expression relative to the Untreated condition, NM cultures exhibited significantly higher expression of pathways related to neuron generation (FDR = 7.92e-10), neuron differentiation (FDR = 16.92e-9), and neuron development (FDR = 9.09e-6), indicating a more robust neurogenic program (Figure 4E). The NM co-culture is also superior to the Untreated and Treated conditions from a mitochondrial function perspective (Figure S3). Untreated neurons while focusing on OXPHOS-based metabolism, showed enrichment of abnormal mitochondrial function pathways and ETC defects. Treatment with microglial mitochondria shifted metabolism away from OXPHOS to glycolysis, bolstering our ECAR data from Figure 2. The NM co-culture was more focused on protein production, with the condition being positively associated with peptide chain elongation and translation. Overall, mitochondrial function was positively effected by treatment, showing enrichment of glycolysis and protein production-related pathways.

However, the NM co-culture allows for a balanced metabolic environment, supports efficient mitochondrial function.

Interestingly, genes associated with negative regulation of nervous system development were also enriched, driven by the differential expression of PTN and ASCL2 (upregulated in Treated) and TLR2 and TREM2 (upregulated in NM). The latter two, known as microglia-associated, likely reflect the broader functional roles of intact microglia beyond mitochondrial support, including their influence on neuroimmune signaling and developmental pruning (Figure 4D).

Collectively, these findings indicate that while mitochondrial transfer alone induces partial maturation and transcriptional shifts in neurons, it does not fully recapitulate the gene expression landscape promoted by direct neuron–microglia co-culture. This suggests that additional microglial-derived factors or sustained physical interaction are necessary to achieve the full spectrum of developmental support.

## Discussion

### Metabolic Reprogramming Reflects Developmental Maturation

Our data demonstrates that the exogenous delivery of microglial mitochondria to neurons induces a temporal shift in metabolic activity that closely resembles developmental transitions observed in human neural maturation (34). At 48 hours post-treatment, we observed a normalization of both oxidative phosphorylation (OXPHOS) and extracellular acidification rate (ECAR) to levels comparable to neuron–microglia (NM) co-cultures. By one week, OXPHOS activity was substantially elevated while ECAR remained unchanged, indicating a durable metabolic reprogramming toward aerobic respiration. This mirrors patterns seen in human stem cell-derived neurons, where a developmental switch from glycolysis to oxidative metabolism is essential for neuronal differentiation, dendritic growth, and synaptic activity (35–38). These results support the view that mitochondrial function is not merely permissive but actively instructive in establishing neuronal metabolic identity.

### Transcriptomic Shifts Support Neurodevelopmental Progression

RNA sequencing revealed that mitochondrial treatment alone is sufficient to trigger broad transcriptional programs associated with nervous system development, including neurogenesis, neuron differentiation, projection development, and both central and peripheral nervous system formation. Genes involved in mitochondrial structure and function—particularly membrane-bound respiratory complexes—were also significantly upregulated. These results align with recent developmental studies implicating mitochondrial biogenesis and cristae remodeling as key drivers of neural lineage specification (34). Specifically, compared to NM conditions, treated neurons still showed enrichment in developmental pathways, including axis elongation and neurogenesis, suggesting that mitochondrial transfer induces a unique and sustained maturation signal.

### Structural and Functional Implications of Mitochondrial Transfer

While metabolic and transcriptional indicators of maturation were strongly enhanced, mitochondrial treatment did not rescue neuronal fragmentation, which remained significantly higher than in NM cultures. This indicates that while mitochondria contribute substantially to the developmental trajectory, full structural support may require additional microglia-derived factors, such as cytokines, extracellular matrix components, or phagocytic signaling. At the protein level, we observed increased expression of MAP2, suggestive of enhanced dendritic architecture (39). However, Syn1 and Tuj1 levels remained unchanged, indicating that synaptogenesis and cytoskeletal integration may be later-stage processes or require broader environmental cues.

### Mitochondria as Drivers of Neuronal Maturation

Collectively, these findings add to a growing body of evidence that mitochondria play a central and active role in neuronal maturation. Beyond energy supply, our data suggest that microglial mitochondria can function as autonomous developmental cues—promoting metabolic reprogramming, initiating neurodevelopmental gene expression, and enhancing structural features of neuronal identity. While the full complexity of neuron–microglia interactions is not replicated by mitochondrial transfer alone, this study provides functional evidence that mitochondria are sufficient to initiate key aspects of the neuronal maturation program. These insights highlight the therapeutic potential of mitochondria-based strategies and underscore their relevance in developmental and regenerative contexts.

### Conclusions and Future Directions

In summary, our findings reveal that microglia and their mitochondria are critical regulators of neuronal maturation and mitochondrial health. These results position microglial mitochondria as key modulators of central nervous system development. Future studies will explore the long-term transcriptomic consequences of microglial mitochondria treatment on naïve neurons and extend our 3D *in vitro* findings to *in vivo* models.

### Limitations

While we provide insight into microglial mitochondria’s potential role in neuronal maturation, it is important to consider the limitations of our study. Firstly, our study lacks multiple genetic backgrounds, which limits our ability to understand individual variability in neurodevelopment and mitochondrial function. In future studies, incorporating multiple donor-derived cell lines will alleviate such limitations and allow for understanding how genetic differences may potentially influence neurodevelopment. Secondly, our study lacks an *in vivo* component. Due to ASD being a behavioral disorder, future studies using mice will be necessary to understand the contribution of microglial mitochondria in neuronal maturation. Lastly, RNA sequencing was conducted only on the 48-hour treated condition, preventing us from capturing potential long-term transcriptomic changes (1wk and beyond). Addressing these limitations in the future will deepen our understanding of microglia mitochondria’s contribution to neurodevelopment.

## Author Contribution

VL and SPS conceived the project. VL and SPS designed and interpreted the experiments and wrote the manuscript. SBSK provided fluorescent mitochondria cell lines. SPS and VL performed scaffold seeding and sample maintenance (media changes). SPS performed all mitochondrial treatments. SPS and VL performed mitochondrial isolation. SPS performed all mitochondrial bioenergetic function assessments. VL performed confocal imaging. SPS and CA conducted Western Blots. SPS performed aspect ratio, mtDNA quantification, colocalization, and neuronal network analysis. SPS prepared silk materials. SPS and VL conducted RNA isolation. SPS, VK, and SBSK performed RNA sequencing analysis.

## Acknowledgements

The authors thank the University of Cincinnati Start-up funds to Volha Liaudanskaya, the University of Cincinnati URC Faculty Scholar Award, NIH 1R21AG085052-01A1, DoD HU00012420075, and the Cincinnati Children’s Hospital Bioimaging and Analysis Facility.

## Methods

### 3D Silk Scaffold Model Preparation

#### Silk Processing and Scaffold Fabrication

Silk scaffolds were prepared from silk fibroin following a previously established protocol (32, 40). Silk cocoons were boiled in a deionized water sodium carbonate solution (0.02□M) for 30□minutes to remove sericin. The silk fibroin fibers were dried overnight in a fume hood. The next day, the silk fibers were dissolved in 9.3□M lithium bromide (Sigma, cat. no. 213225) by immersion for 4□h at 60□°C. Following the dissolving of the silk, dialysis of the silk in deionized water at room temperature was performed for 3 days in dialysis tubing to remove the lithium bromide from the silk (total of 6 water changes).

The resulting silk solution was centrifuged twice at 9,000□rpm for 20□minutes, and a 100µm strainer was used to filter out the remaining debris. Then, 500µL of the silk solution was dried in a small weight boat at 60□°C overnight to determine the weight/volume (w/v) concentration. The silk solution concentration was adjusted to 6□mg/mL and combined in a 10□cm dish with 400 to 500 µm sodium chloride (Sigma, cat. no. 71382) at a ratio of 1:2 (v/w). The silk solution was then left for 2 days at room temperature to trigger pore formation of the 3D silk sponge; the solution was then incubated for one hour at 60□°C. After separating the sponge from the dish, residual sodium chloride was removed by dialysis in deionized water for 2 days at room temperature with 6 water changes total (3 times a day).

To create the scaffolds for experimental purposes, the sponges were cut into individual donut-shaped scaffolds using biopsy punches (Integra) with a 6□mm OD (outer diameter) and 2□mm ID (inner diameter). The scaffolds were trimmed to a height of 1.5□mm using scissors. Following the shaping of the scaffolds, they were then autoclaved in deionized water for 20□minutes with the liquid cycle, cooled to room temperature, and then processed for experiments (process described below).

#### Coating with ECM

Extracellular matrix coating of scaffolds was done using previously generated protocols (32). In brief, scaffolds were coated with 10□µg/mL poly-ornithine (PLO) (Sigma, cat. No. P4957) and 5□µg/mL laminin (Fisher, cat. no. 501003381) to allow for proper cell adhesion. Scaffolds were placed into a 6-well plate (maximum of 50 per well) using tweezers and incubated overnight at 37□°C in 7□ml of a 10ug/ml PLO coating solution in distilled water. The next day, the PLO was aspirated and washed three times with PBS. The scaffolds were then incubated overnight at 4□°C in 0.5□mg/mL 100μL laminin stock aliquots with Dulbecco’s Modified Eagle Medium/Nutrient Mixture (DMEM F12) phenol red-free media in a 1:100 ratio and then the scaffolds were retained in the laminin solution. Before use, the scaffolds were rinsed with PBS, followed by adding 5mL of media of interest.

#### Cell Culture

##### Fluorescent Mitochondria Cell Lines (mtDsRed2 hiNSCs and mtEBFP2 Microglia)

We used previously generated in our lab mt-dsRED2 hiNSCs and mt-BFP2 microglia following the established protocol (41). In brief, hiNSCs and HMC3 human microglia cells were transduced with hSYN-mito-dsRED2 plasmid (Addgene, 173069) for hiNSCs and a custom plasmid used VectorBuilder to implement a double COX8A promoter and EBFP2 into the plasmid backbone for HMC3s (41, 42).

##### Mouse Embryonic Fibroblasts (MEFs)

Following ATCC protocols, MEFs were used as a feeder layer for induced human neuronal stem cells (ihNSCs). The cells were seeded on a 15cm^2^ gelatin-coated dish (0.1% gelatin for 20 minutes: Sigma-Aldrich SF008). MEFs were used at passages 3-4. DMEM supplemented with 10% Fetal Bovine Serum (FBS), and 1% Antibiotic-antimycotic (Anti-Anti) were used to maintain the MEFs. The media was changed every 3 days.

Once the MEFs reach 100% confluency, they are then inactivated with 10□µg/mL Mitomycin C (Sigma, M4287) for 3Lhours, followed by three washes with Phosphate Buffered Saline (PBS). After the last PBS wash, media was added to the plates, for hiNSC seeding.

##### mt-DsRed2 Induced Human Neural Stem Cells (hiNSCs)

mt-dsRED2 hiNSCs were handled and maintained following previously established protocols (41). The media was composed of KnockOut DMEM supplemented with 1% GlutaMax, 20% KnockOut Serum Replacement, 1% Anti-Anti, 0.2% β-mercaptoethanol (Invitrogen), and 800μL of 10□µg/mL basic fibroblast growth factor (Invitrogen) added proportionally to aliquots of media upon use. Media was changed every other day. mt-dsRED2 hiNSCs were expanded and cultured on top of inactivated mouse embryonic fibroblasts. When cells reached 70–80% confluency, they were lifted from the plates via incubation with TrypLE solution for 1□min at 37□°C, followed by inactivation with mt-dsRED2 hiNSCs media. Cells were then pelleted by centrifugation at 3000□rpm for 2□minutes. For further expansion, pellets were gently mechanically disrupted with a 5□ml pipette, and the colony suspension was transferred to Matrigel-coated dish (1:50 dilution of Matrigel in phenol red free DMEM F12 for 1 hour), mt-dsRED2 hiNSCs colonies were replated into the Matrigel coated dishes following 1 to 20 15cm^2^ dishes ratio. For 3D seeding, the pellet was rigorously pipetted to obtain single-cell suspension along with centrifugation at 1000rpm for 5 minutes.

##### mt-EBFP2 HMC3 Microglia Cell Line

The transduced mt-EBFP2 HMC3 microglia cell line was maintained following established ATCC protocols (41). Fluorescent mitochondria microglia were expanded in 15 cm^2^ dishes in EMEM media supplemented with 10% FBS, and 1% Anti-Anti. The media was changed every 3 days. When cells reached 70-80% confluency, they were lifted from plates with 0.25% Trypsin/EDTA for 3□min at 37□°C. After adding complete media to the mt-EBFP2 microglia, cells were pelleted with centrifugation at 1000□rpm for 5□minutes. Cells were replated at 300,000 cells/cm^2^.

##### 3D *in vitro* Human Brain Tissue Model Fabrication

The system for generating 3D human *in vitro* brain tissue model was done via an earlier published protocol (32). Once mt-DsRed2 hiNSCs and mt-EBFP2 HMC3s reached nearly 100% confluency, they were collected as described in the individual culture methods and combined to achieve the following ratio: 2:0.1 million neurons and microglia, respectively. The same ratio was used for monocultures for the respective cell types. Immediately before seeding the mono- or co-culture cell solution, PLO- and laminin-coated silk scaffolds were placed in 96-well plates with tweezers and an aspirator line to remove excess liquid. Calculations were performed so that each 40μL of cell suspension contained 2:0.1 million mt-DsRed2 neurons and mt-EBFP2 microglia. Then 40μL of the cell suspension was pipetted onto the semi-dried scaffolds and incubated for 30□minutes at 37□°C to allow the cells to attach. After this, 150μL of complete neurobasal media was added to each well and then placed in an incubator at 37□°C with 5% CO_2_ in a humidified atmosphere for 24 hours to allow cells to sufficiently attach to silk scaffolds. Complete neurobasal media (CNB) was composed of Neurobasal medium supplemented with 2% B-27, 1% Anti-Anti, 1% Glutamax, and 1% astrocyte growth factors.

The following day, the cell-seeded scaffolds were transferred to new 96-well plates using sterile tweezers. To ensure a full 3D environment, each scaffold was embedded in100μL of collagen type I solution with complete neurobasal media, 10x PBS, and a pH adjusted to 7.0-7.2 with NaOH and then incubated for 30□minutes at 37□°C to allow the collagen gel to crosslink. Next, 150μL of complete neurobasal media was added to all scaffolds and incubated for 24 hours at 37□°C. The next day, the brain-like tissues were moved into 48-well plates with 1□ml of complete neurobasal media in each well. Half of the media changes were done every fourth day until the tissues were mature enough for experimental analysis (dense neuronal network, and spontaneous neuronal activity or 5-6 weeks after seeding (43).

#### Mitochondria Isolation, Treatment, and Analysis

##### Microglia Mitochondria Isolation and Treatment of 3D *in vitro* Brain Cultures

###### Intracellular Isolation

Mitochondria isolation was conducted using the established protocol from the Thermofisher Scientific Mitochondria Isolation Kit for Cultured Cells (Thermo Fisher: 89874). In brief, 6wk old scaffolds were transferred into 2ml Eppendorf tubes and serial treated with supplemented reagents A-C, followed by mechanical breakage of the scaffold, and 5 minutes max speed vortexing. Next, the large pieces of silk debris were removed from the Eppendorf tubes, followed by removal of small ECM, cell, and silk debris after 10 minutes centrifugation at 1000 x g (4°C). The mitochondria fraction was pelleted by 25 minutes of centrifugation at 12000 x g (4°C). Isolated mitochondria were either used for treatment or analysis. The residual protein-rich supernatant was stored in 2mL Eppendorf tubes at −80°C for Western Blot analysis.

###### Extracellular Isolation

The media from 3D in vitro cultures was collected into the 1.5□ml Eppendorf tubes and centrifuged for 10□minutes at 1000xg to pellet the cells. The supernatant was centrifuged at 12,000x g for 25□minutes to collect a mitochondria-enriched fraction. The residual mitochondria-free media after centrifugation was stored at −80°C for later analysis.

###### Treatment of Naïve Neurons

Mitochondria enriched CNB - immediately before the treatment, the pelleted microglia mitochondria was reconstituted in 1ml of CNB media, and this solution was then used for the treatment protocol. In brief, to deliver the mitochondria-enriched CNB, the entire media volume was removed from the naïve neuronal cultures (48-well plate - 1mL) and replaced with 1mL of mitochondria-enriched CNB for the “Treated” conditions. “Untreated” and “NM” conditions also received a full media change to maintain consistency within the experiment.

### Mitochondria Bioenergetic Analysis via Seahorse Assay

The mitochondria’s bioenergetic function was measured using Seahorse Real-Time ATP Rate Assay following the manufacturer’s instructions with minor optimizations for our system. In brief, 12-18 hours prior to running the Seahorse Real-Time ATP Rate Assay, the sensors from the Extracellular Flux Pack (Agilent Technologies: 103792-100) were hydrated with 200µL of sterile water the day before the assay and stored in a 37°C non-CO_2_ incubator overnight. The day of the assay, the sterile water was removed and 200µL of Seahorse XF Calibrant was added to the utility plate for at least 2-4 hours. The assay medium was prepared with 50mL of phenol-red free DMEM F12, 10mM of XF glucose, 1mM of XF pyruvate, and 2mM of XF glutamine. Stock compounds were made on the day of the assay with resulting stock concentrations of 150 µM for Oligomycin and 50 µM for Rotenone + Antimycin A. These stock solutions were then diluted in prepared media (1:10).

Following the preparation of assay compounds and Seahorse media, mitochondria were isolated from the 3D silk scaffolds following the procedure above at select time points (24hrs, 48hrs, and 1wk post-treatment). Once the mitochondria (intracellular and extracellular) were isolated, it was then reconstituted in 180µL of seahorse media and added to the Cell Culture Microplate. All empty wells were filled with 180µL of seahorse media to be used as a negative control and blanks for the assay. Using the established protocol from Agilent Technologies for the Standard Assay, 20µL of the final prepared Oligomycin is added into Port A and 22µL of Rotenone + Antimycin A into Port B of the Sensor Cartridge.

After the loading of the mitochondria and assay compounds, the mitochondria were analyzed via the Seahorse XFe96 Analyzer using the “Real-Time ATP Rate Assay” template to measure Oxygen Consumption Rate (OCR) and Extracellular Acidification Rate (ECAR). Data collected was used for further analysis of mitochondrial function. Succeeding the Real-Time ATP Rate Assay, 20µL of the mitochondria-rich media is stored at −20°C for mtDNA quantification. The remaining 160µL is centrifuged at 12,000xg for 20 minutes and reconstituted in 60µL of RIPA (supplemented with protease and phosphatase inhibitors) buffer for storage and later analysis.

### Mitochondria Transfer and Progression of Neuronal Maturation

#### Immunofluorescence Staining and Analysis

PBS solution supplemented with 4% sucrose and 4% paraformaldehyde (PFA, Electron Microscopy Sciences), was used to fix the scaffolds at the selected (indicated) timepoints. After fixing, the tissues were washed four times with PBS and then permeabilized for 1 hour with a permeabilization solution composed of 0.2% Triton X-100 and 4% goat serum. Using the permeabilization solution, primary antibodies were diluted (anti-Tuj1: ab78078 mouse; 1:1000) and added to the scaffolds. The scaffolds were incubated in the primary antibody at 4□°C overnight on a rocker, then washed with gentle shaking in PBS four times (10□minutes each wash). After washing, tissues were incubated on a shaker with secondary antibodies diluted in PBS for 1□hour at room temperature (goat anti-mouse Alexa 647: A11126; 1:500). Four additional PBS washes were used to dispose of any unbound antibodies. For all figures, fluorescent image stacks of the stained scaffolds were acquired on a Nikon A1 inverted LUNV confocal microscope (Nikon, Plan Apo λ 20x, 1024×1024, each maximum projection had 50 steps). Images in the figures represented maximum intensity projection and were collected with the same PMT gain settings and laser power between all experiments. All scaffolds were imaged with the EBFP2 line blind to avoid bias. Tuj1 neuronal network density was analyzed using a custom MATLAB code (44).

### Cell-Specific Mitochondrial Aspect Ratio Analysis

In brief, the ratio of mitochondria length versus width (aspect ratio) was measured using a publicly available macro from NIH ImageJ software (45). Max projection images of fluorescent mitochondria acquired with Nikon A1 inverted LUNV confocal microscope (1024×1024 pixels, 20x objective) were analyzed via the macro. The measured aspect ratio for each mitochondrion was plotted as a distribution graph for comparison with other groups.

### Mitochondrial DNA Quantification

Using previously established protocols with minor optimizations for our system [Invitrogen: P11496]. Previously prepared samples from the Seahorse Assay were thawed on a shaker and buffer solutions were prepared (1X TE buffer and 0.2% Triton X). While waiting for samples to thaw, standards for the standard curve were prepared in a 48-well plate with 1X TE buffer and DNA working solution (working solution: 1:50 dilution of DNA Lambda in 1X TE buffer). The final concentrations for the standards are as follows, 2000ng/mL, 1000ng/mL, 500ng/mL, 200ng/mL, 100ng/mL, 50ng/mL, 20ng/mL, and 0ng/mL. Once the samples are thawed, dilute your sample with 1X TE buffer and 0.2% Triton X (1:5 dilution of sample) for a total sample volume of 100μL. Once the samples are prepared, they are transferred to a black 96-well plate. The PicoGreen solution was prepared so there is 100μL per sample and standard (1:200 dilution in 1X TE buffer).

Prior to the addition of the PicoGreen Solution, the plate reader was set to fluorescent excitation = 485nm, emission = 538, and shake for 5 seconds. Once the plate reader is set up. In a dim room, add 100μL of PicoGreen Solution to the wells with samples and standards, then read the plate, continuing reading until the reaction is completed. For analysis, a standard curve was generated from the standards on the plate. Followed by dilutions and application of standard equation to generate mtDNA content per sample.

### Mitochondria Colocalization Analysis

Colocalization analysis was completed using the JACoP plugin from the NIH ImageJ software (46). Images were acquired with Nikon A1 inverted LUNV confocal microscope (1024×1024 pixels, 20x objective) were analyzed via the macro with the Otsu thresholding method. Spearman’s correlation coefficients were plotted by condition for comparison with other groups and/or timepoints.

### Western Blot

For whole scaffolds, 1X RIPA lysis buffer (with protease and phosphatase inhibitors) was added to extract cellular protein lysates. Scaffolds were subsequently sonicated at 20% amplitude for 20 pulses (1-second pulse on, followed by 1-second off). Sonicated samples were then diluted at a 3:1 ratio with a sample buffer (9:1 Laemmli buffer to β-mercaptoethanol). Samples containing isolated mitochondria (intracellular and/or extracellular) or total protein fraction were obtained during the Seahorse Assay and post-processing. For analysis, 15µL of sample were mixed with 5 µL of sample buffer (15-well gel), while post-Seahorse Assay protein samples were prepared with 45 µL of sample and with 15 µL of sample buffer (10-well gel). Prepared samples were heated at 95°C for 5 minutes prior to loading onto pre-cast gels for electrophoresis. Electrophoresis was conducted at 150 V for approximately 50 minutes to 1 hour. Gels were imaged using the Stain-Free preset on a Bio-Rad imager to obtain total protein volume per lane.

If sample numbers exceeded the well capacity of a single gel, samples were split across two gels. To ensure consistency in detection, gels were sectioned at the same molecular weight markers (75 kDa and 20 kDa) and transferred onto a polyvinylidene fluoride (PVDF) membrane. Protein transfer from gels to PVDF membranes was performed using a semi-dry transfer system at 2.5 V and 1.6 A for 4 minutes (for total protein samples) or 3 minutes (for mitochondrial samples). Membranes were imaged using stain-free blot imaging to assess total protein content, facilitating subsequent normalization.

Membranes were blocked for 5 minutes using Everything Blocking Buffer (Bio-Rad, 12010020) and then were incubated with primary antibodies diluted in Everything Blocking Buffer at 4°C overnight (primaries listed in Table 1) overnight at 4°C on a rocker. After primary incubation, membranes were washed with 1X Tris-Buffered Saline with Tween 20 (TBST) (2 instant washes, followed by 3, 10-minute washes) to remove excess primary antibody. Membranes were then incubated with secondary antibodies (1:10,000 dilution in Blocking Buffer, Cell Signal Technologies, HRP – conjugated: anti-rabbit IgG 7074P2; anti-mouse IgG 7076S) for 1 hour at 4°C, followed by the same wash sequence. For signal detection, membranes were exposed to an ultra-sensitive enhanced chemiluminescent (ECL) substrate for 5 minutes before chemiluminescence imaging. Following imaging, membranes were stripped with a western blot stripping buffer at room temperature for 30 minutes, followed by the same TBST wash sequence, to enable repeated antibody staining for multiple target proteins. The images were analyzed using ImageJ Software.

**Table 1:** Primary Antibodies.

| Primary Antibody | Animal | Dilution | Product Code |
| --- | --- | --- | --- |
| Anti-pDRP1 | Rabbit | 1:1000 | Cell Signaling – 3455S |
| Anti-DNM1L | Mouse | 1:1000 | Invitrogen – MA5-26255 |
| Anti- Synapsin 1 | Rabbit | 1:1000 | Abcam – AB64581 |
| Anti-MAP2 | Rabbit | 1:1000 | Abcam – AB32454 |
| Anti-CX3CL1 | Mouse | 1:1000 | Invitrogen – MA1-031 |
| Anti-MTCO1 | Mouse | 1:1000 | Invitrogen - 459600 |
| Anti-VDAC1 | Rabbit | 1:1000 | Cell Signaling - 4661S |
| Anti-Tuj1 | Mouse | 1:1000 | Abcam - AB78078 |
| Anti-TOMM20 | Rabbit | 1:1000 | Invitrogen - MA5-24859 |

### RNA Isolation Protocol

Scaffolds were collected and stored at −80°C. Upon isolation, samples were thawed on ice and incubated with 600□µL of Lysis Buffer from RNeasy Kit (Qiagen, Catalog No. 74106). During incubation, samples were cut up using small scissors and sonicated to break up the scaffold (Amplitude – 20%, Time – 5 seconds, 1 second on 1 second off). After homogenization, the samples continued to sit in a lysis buffer on ice for approximately 20□minutes. The lysed samples were placed onto QIAshredder Mini Spin (Qiagen) columns to remove pieces of silk and spun at 15,000□rpm for 2□minutes. An equal amount of 70% ethanol was added to the supernatant and mixed thoroughly. The ethanol-containing mixture was spun in two rounds through the RNeasy Mini Spin column (Qiagen) at 10,000□rpm for 1□minute each; the flow through was discarded. In a new collection tube, 350µL of RNA wash from RNeasy Mini Kit (Qiagen) added to the spin column and incubated for 5 minutes. The sample was added to the RNeasy column and spun at the same setting for 1□minute; the flow through was discarded. After the first RNA Wash, TURBO DNase treatment was prepared and completed following manufacturer’s instructions (Invitrogen, cat. no. P5537155). After the final spin at 10,000x g for 1.5 minutes, a second and third round of RNA was performed as well as a final spin to ensure the column was dry. The column was placed into a 1.5□mL microcentrifuge tube, and 24µL of RNase-free water (Fisher Scientific) was added to the center of the column, incubated for 10 minutes, and spun at 10,000□rpm for 1□minute (two times). Samples were then read on the Tecan Infinite M Plex NanoQuant Plate™ and stored at −80°C before sequencing.

Azenta Life Sciences completed sequencing, data was analyzed using Python comparing NM to Treated and Treated to Untreated conditions (p-value < 0.05, log_2_(fold change) = 0.2). Differentially expressed genes were then compared to the MitoCarta 3.0 to delve into potential bioenergetic pathways (33). Heatmaps and volcano plots were generated with matplotlib Python package.

### Statistical Analysis

Statistical analysis was performed between and within experimental groups using GraphPad Prism software. Two-tailed t-tests were used to compare values within an experimental group and between two experimental groups. Mann-Whitney U and Welch’s t-tests were used when necessary. One- or two-way ANOVA analysis of variance was used to compare multiple groups. Tukey’s post hoc tests were used to assess computed significant differences between experimental and control groups when the data was normal. Kruskal-Wallis was used with Dunn’s post-hoc test when data was nonparametric. Repeated measure ANOVA was used to compare control and experimental conditions, when necessary, followed by Tukey post-hoc test. Any p-value less than 0.05 was considered statistically significant. Each experiment was repeated at least two times; technical replicates (2-3, depending on the experiment) were used for every assay. Data was represented by mean and standard error of the mean (SEM) of each group. Power analysis was completed at a power of 80% with an α value of 0.05, leading to n = 3 for sham and n = 4 for experimental condition.

**S1:**
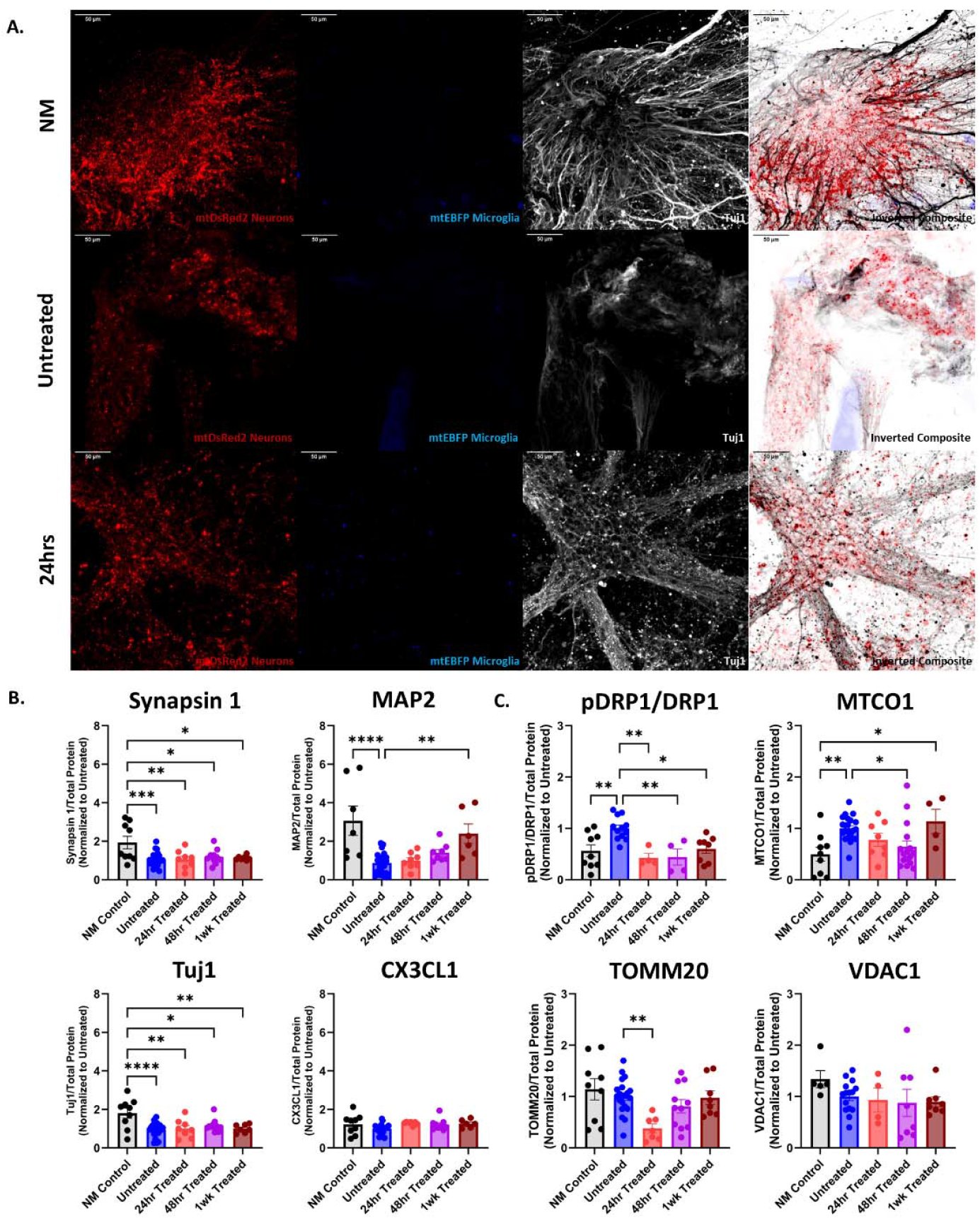
24hr Treatment – Maturation and Mitochondrial Health: **A**. Representative images of NM, Untreated, and 24hrs Treated. mtDsRed2 neurons and mtEBFP2 microglia. Tuj1 staining for neuronal networks. Nikon A1 inverted LUNV confocal microscope 20x objective 1024×1024 pixels with 50-steps Max projection. Scale bar: 50µm. **B**. Neuronal maturation markers MAP2, CX3CL1, Synapsin 1, and Tuj1 over time western blots normalized to total protein concentration per lane and to Untreated condition. **C**. Mitochondria health markers, pDRP1/DRP1, TOMM20, VDAC1, and MTCO1 intracellular mitochondria western blots normalized to total protein concentration per lane and to Untreated condition. n = 3 for NM and Untreated. n = 4 for 24hr Treated. Mean ± SEM. One-way ANOVA with Tukey post-hoc test and α = 0.05. ROUT outlier analysis method. Analysis completed in GraphPad Prism. n = 3 for NM and Untreated. Each timepoint was replicated at least twice.

**S2:**
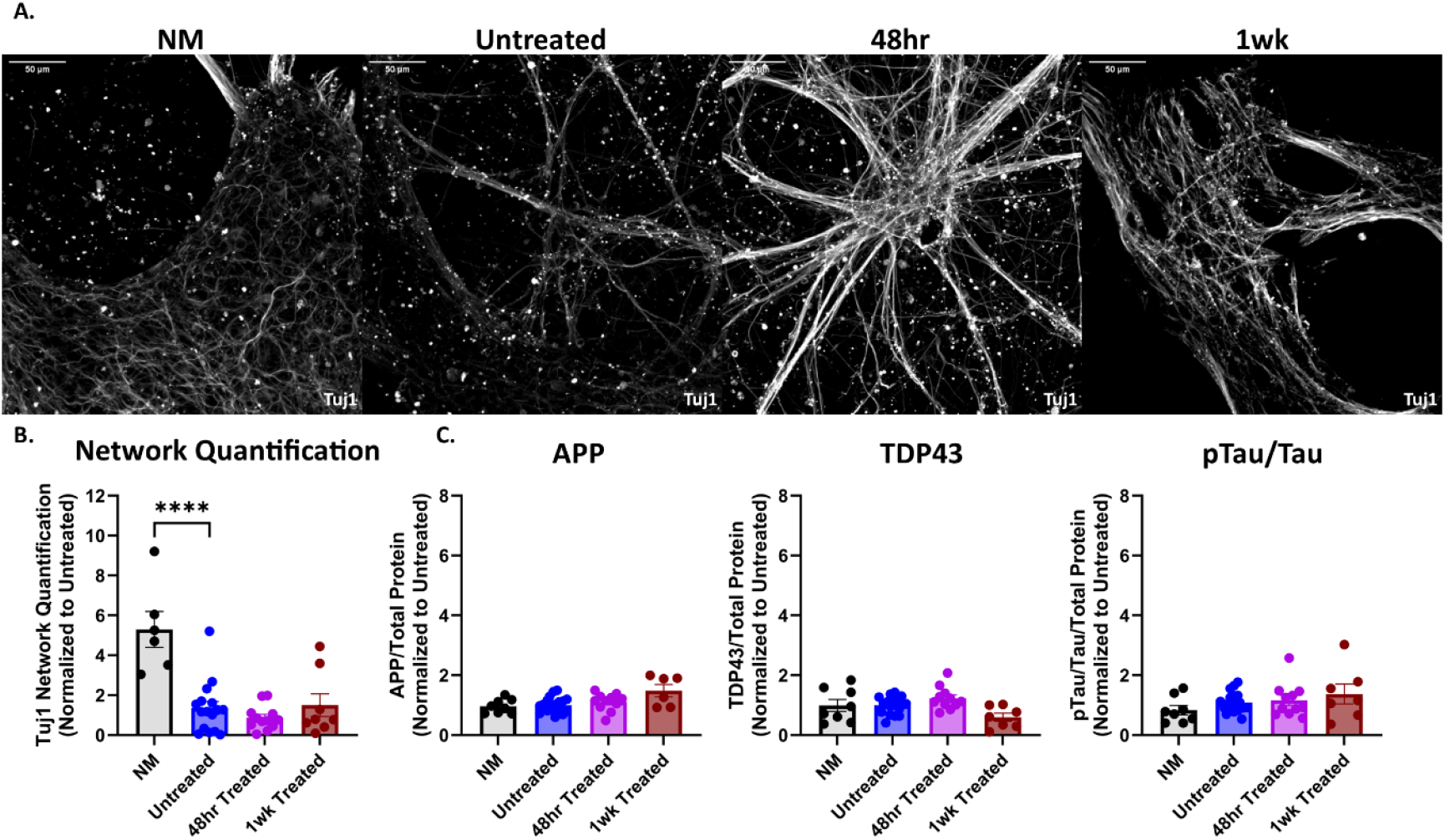
Treatment and Neurodegeneration: **A**. Representative images of NM, Untreated, 48hrs, and 1wk Treated. Tuj1 staining for neuronal networks. Nikon A1 inverted LUNV confocal microscope 20x objective 1024×1024 pixels with 50-steps Max projection. Scale bar: 50µm. **B**. Neural network quantification with custom MATLAB code comparing NM, Untreated, 48hrs, and 1wk Treated (44). **C**. Neurodegeneration markers, APP, TDP43, and pTau/Tau western blots normalized to total protein concentration per lane and to Untreated condition. n = 3 for NM and Untreated. n = 4 for 48hr and 1wk Treated. Mean ± SEM. One-way ANOVA with Tukey post-hoc test and α = 0.05. ROUT outlier analysis method. Analysis completed in GraphPad Prism. n = 3 for NM and Untreated. Each timepoint was replicated at least twice.

**S3:**
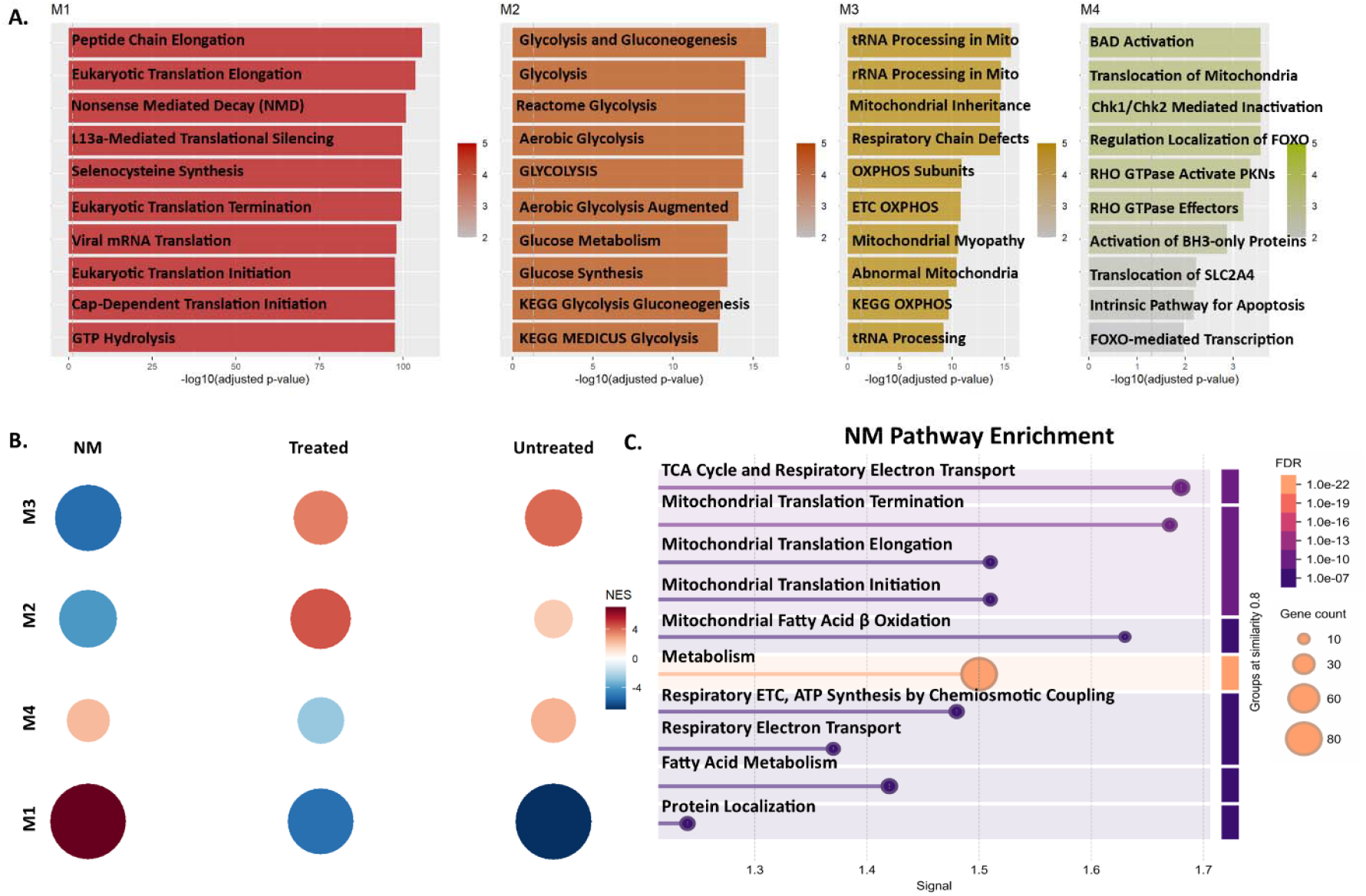
CEMiTool Pathway Enrichment: **A & B**. Gene set enrichment analysis demonstrating 5 modules with M1 – Protein Production (positively associated with NM), M2 – Glycolysis (positively associated with Treated), M3 – OXPHOS and Mitochondria Dysfunction (negatively associated with NM, positive with Untreated), and M4 – Mitochondrial Motility (positively associated with Untreated). **C**. NM mitochondrial pathway enrichment genes from Mito Carta 3.0 in NM. Log_2_(Fold Change) = 0.5, adjusted p-value of < 0.05, medium confidence interaction score (0.4), and FDR ≤ 0.05 (33).

